# Excitable dynamics of ParA during bacterial cell division

**DOI:** 10.64898/2025.12.16.694648

**Authors:** Jerome Rech, Oliver Valera, Longhua Hu, Celine Mathieu-Demaziere, Tanya Nesterova, Jean-Yves Bouet, Jian Liu

## Abstract

ParABS system is a conserved machinery that drives the partitions of genome and other large intracellular cargos in bacteria. It works as a ParA concentration-based Brownian ratchet, which ensures the partition fidelity by operating at the critical point in its parameter space and localizing the ParA nearby to buffer against the large fluctuations in the ParA intracellular concentration. Despite these progresses, our understanding of the ParABS-driven partition mechanism remains incomplete. It is not understood what causes the large cell-to-cell fluctuation in the ParA intracellular concentration and why the bacterium cannot minimize this noise to better control the partition fidelity. We present experimental evidence with low-copy-number F-plasmids in *E. coli* that ParA robustly oscillates from pole to pole along the nucleoid length, underlying the observed large fluctuations in ParA intracellular concentration. Our theory-experiment synergy demonstrates that the ParA oscillation hinges on the nucleoid dimension and the ParB-mediated negative feedback with ParA in nucleoid binding. We suggest that the low-copy-number plasmid manages its partition fidelity with “autonomy”: The plasmid harnesses the same ParB-mediated negative feedback – that dissociates the ParA from the nucleoid underneath the plasmid – to maintain the nucleoid-bound ParA at a proper level further away. And the ParA oscillation is reminiscent of this underlying feedback mechanism, which adapts the partition fidelity to the nucleoid growth. Our work may provide an overarching framework of intracellular partitioning in bacteria.

**Significance Statement:** ParABS system is a conserved partition machinery for genome and other large cargos inside bacteria and works as a ParA concentration-based Brownian ratchet. It operates at the critical point in parameter space for sensitive adaptation and localizes the ParA nearby to buffer against the large cell-to-cell ParA concentration variations evidenced in bacteria. Why cannot the bacterium minimize this noise to better control the partition fidelity? Hereby, we showed that the large cell-to-cell ParA concentration variations stem from the ParA pole-to-pole oscillations along the nucleoid length and embodies an automony strategy that the intracellular cargo controls the nucleoid-bound ParA at a proper level to ensures its partition fidelity. Our work may provide an overarching framework of intracellular partitioning in bacteria.

## Introduction

ParABS system is a highly conserved machinery across different bacteria that partitions the low-copy plasmids, chromosomes, and other large intracellular cargos by driving directed motilities (1–4). In this tripartite machinery, ParA is an ATPase that binds to the nucleoid nonspecifically; ParB is the adaptor protein that interacts with the nucleoid-bound ParA on one hand and connects to the ParS or alike of the intracellular cargo on the other hand (2, 3). Despite its simplicity of the tripartite interplay between ParA, ParB, and ParS, how the ParABS system underlies the partition process remains not well-understood. Importantly, unlike the linear stepper motor proteins in eukaryotes, the ATP hydrolysis of ParA is demonstrated to only marginally impact the partition process (5). This brings up the fundamental question of how to drive directed motilities without much active energy inputs.

Recent finding from us and others’ have reached some consensus on the ParABS-mediated partition mechanism, although may varied in detail (2, 6–19). Collectively, the current work suggested that instead of forming filaments, the ParABS drives the partition via a ParA concentration-based Brownian ratchet mechanism (7, 9–11, 15, 17). Hereby, the intracellular cargo-bound ParB “kicks” out the nucleoid-bound ParA; this creates a ParA concentration gradient that rectifies the random motions of the cargo into the directional movement. Our work further identified two important features of the ParABS system (15). First, this Brownian ratcheting works at a tipping point in the parameter space (aka “critical point”), which sensitively adapts the separation between the replicated plasmids to the nucleoid length. This way, when there are the two replicated plasmid foci inside the cell, they partition by segregating ∼ half of the nucleoid length apart; and when there are more than two foci, then they are equidistant along the nucleoid length (19–27). This can also explain the robust positioning of other intracellular cargoes, such as carboxysomes (28). Second, the low-copy-number plasmid localizes the ParA nearby, buffering against the large cell-to-cell fluctuations in the ParA concentration (> 10 folds) evidenced in experiments (15). Crucially, the combination of these two features ensures the partition fidelity of low-copy-number plasmids (2).

While these findings have deepened our understanding of the ParABS-driven partition, they raise fundamental mechanistic questions. What underlies the large cell-to-cell variability in ParA concentration, given that both *parA* and *parB* are autoregulated genes (29, 30)? And why does the cell tolerate such fluctuations instead of minimizing them to ensure optimal partition fidelity? Clearly, our current understanding of ParABS partition mechanism remains incomplete.

Zooming out into a broader perspective, dissecting how the ParABS system works will not only define the fundamental principles of the partition processes in bacteria, but also help shed light on how the similar molecular machineries have adapts to the different processes among the prokaryotic world throughout the evolution. Notably, MinCDE system, closely related to ParABS system oscillates along the cell length *in vivo* to regulate the septum positioning bacterial cell division (31), and displays a variety of fascinating excitable dynamics *in vitro* (31, 32). The peculiar features in such excitable dynamics expose the inner working of the intricate feedback between molecular constituents, constrain the possible underlying mechanisms, and help unravel the molecular logic of the feedback control (33, 34). Further, while the ParABS system in bacteria does not form cytoskeletons under physiological conditions (6–8, 11, 14, 35), it performs the similar function of its eukaryote cytoskeletons, such as actin. Interestingly, actin machinery is reported to display the oscillation and wave propagation (e.g., (36–38)) and curvature-mediated membrane waves (39) across many different cell types. In this perspective, excitable dynamics may provide a vista point of view that enables us to appreciate the multi-layered feedback regulation inside cell.

In this work, we set out to determine whether and how ParABS system may harness excitable dynamics for the partition process. We present evidence that like the MinCDE system and actin, ParA exhibits remarkable excitable dynamics inside bacteria. ParA oscillates from pole to pole along the nucleoid length, which is robust against a variety of perturbations, does not tightly couple to the directional movements of the low-copy plasmids nor the nucleoid segregation, and introduces the large cell-to-cell variations in ParA concentration. Importantly, combining theoretical modeling and experimental testing we further demonstrate that ParA oscillation relies on the dimension of nucleoid and the ParB-mediated negative feedback with ParA in nucleoid binding. Together, our work suggests that the low-copy-number plasmid manages its own partition in “autonomy”: The plasmid not only builds up the partition complex (PC) by condensing the ParB molecules around *parS* but also controls its own migratory path on the nucleoid. The latter includes localizing some ParA nearby the PC and regulating the stoichiometry of the free ParA and ParB molecules away from the PC. This way, the PC harnesses the same ParB-mediated negative feedback – that dissociates the ParA from its underneath – to keep the nucleoid-bound ParA far away at a proper level. Moreover, the ParA pole-to-pole oscillation is reminiscent of this underlying feedback mechanism, which adapts the partition fidelity to the nucleoid growth. Given that ParA similarly oscillates during the chromosome partitioning in *B. subtilis* (40) and *V. cholerae* (41) and the carboxysome positioning in cyanobacteria (28), our work may provide an overarching framework to understand the fundamental principles of intracellular partitioning and positioning in bacteria.

## Results

### ParA_F_ displays pole-to-pole oscillation without tight coupling to nucleoid constriction nor PC movements

We monitored the dynamics of ParA from the F-plasmids in *E. coli* (**Fig. 1A**). Throughout this study, the subscript “F” denotes the components of the ParAB*S* partition machinery for the plasmid F (ParA_F_, ParB_F_ and *parS*_F_). ParA_F_ exhibited a robust pole-to-pole oscillation along the length of the nucleoid with a period of 5-15 min (**Figs. 1B, C**). To our surprise, the ParA_F_ oscillation remains largely insensitive to a wide range of perturbations, including variations in growth medium, temperature, transcription, inhibition by rifampicin, or stationary phase (**Fig. 1D**). Importantly, ParA_F_ oscillations persisted throughout the cell cycle, regardless of whether the nucleoid divides: the ParA_F_ oscillation occurs on the nucleoid that is coherent without any discernible sign of constriction and persists after the nucleoid replicates and segregates until the cell divides. The oscillation’s insensitivity to nucleoid segregation was particularly evident in stationary-phase cells, where neither the cell nor the nucleoid divided, yet ParA_F_ oscillations remained active (**Fig. 1E**). Furthermore, the ParA_F_ pole-to-pole oscillation was not tightly correlate with the directional movements of PC as shown with the dual-labelling of ParA_F_ and ParB_F_ (**Figs. 1F and S1**), consistent with previous observations showing that the PC of plasmid F remained mostly anchored to the patches of immobilized ParA_F_ on the dense regions of the nucleoid (14, 15).

**Fig. 1.**
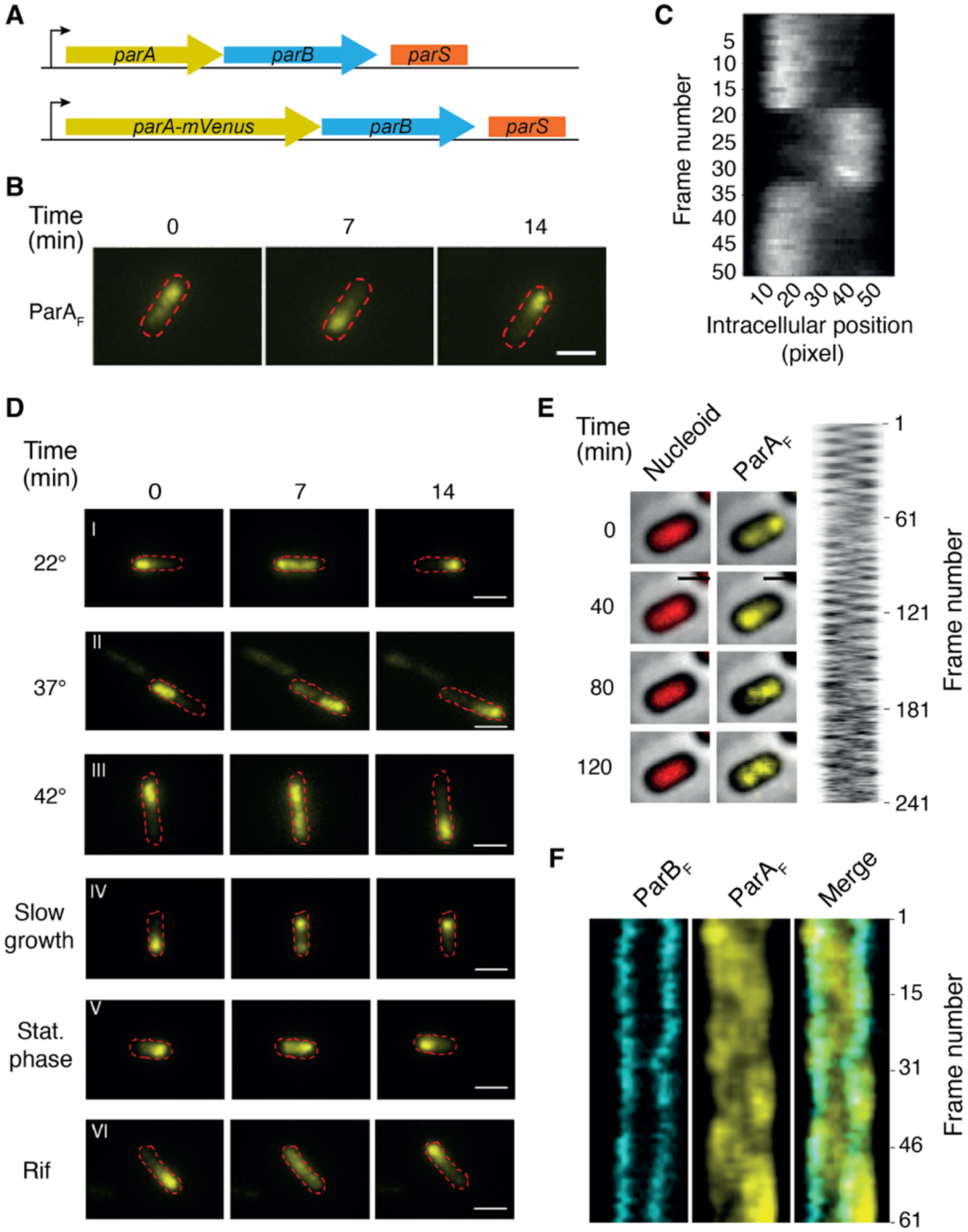
Experimental characterization of ParA_F_ oscillation in *E. coli*. A) Schematics of the genetic constructs. B) Representative live-cell microscopy images displaying ParA_F_ oscillation along the length of the cell. C) Representative kymograph of ParA_F_ oscillation, with images acquired every 30 sec. D) ParA_F_ oscillation is robust to various perturbations, including changes in growth medium, temperature, and transcription arrest. E) ParA_F_ from stationary phase cell oscillates without discernible constriction of the nucleoid and continues after the nucleoid splits. A late stationary phase *E. coli* culture carrying the mini-F pJYB243 was imaged every 30 sec over 120 min in bright field, red, and yellow channels to observe the cell contour, the Hu-mCherry labelled nucleoid and ParA_F_-mVenus, respectively. *Left*: serial images of combined bright field with the red and yellow channel at 0, 40, 80, and 120 min showing that the cell is not growing and displays a homogeneous nucleoid over which ParA_F_ oscillates from both edges. *Right*: Kymograph from the same cell as in the left panel, displaying ParA_F_-mVenus fluorescence over time. F) ParA_F_ oscillation does not correlate with directional movement of PC. Stationary phase culture of *E. coli* carrying pJYB243 expressing *parA*_F_-mVenus and *parB*_F_-mTurquoise2 were imaged every 30 sec for 30 min. Kymographs in the mTurquoise2 (*left*), mVenus (*central*), and merged (*right*) channels are shown. Scale bar: 1 µm (B, D-F).

These observations thus contrast a recent report claiming that ParA_F_ oscillation waves are triggered by nucleoid splitting and depend on PC movement (42). To understand these differences, we conducted further experiments and found that exposure to blue light (438 nm) induced cell growth arrest with most *E. coli* cells displaying apparently segregated nucleoids (**Fig. S2**): *i.e.*, these cells spend most of their time with segregated nucleoids. We showed that under these conditions ParA_F_ oscillated between the segregated nucleoids within the same cell (**Fig. S2**) as reported in (42). However, we note that just because ParA_F_ oscillation was observed under these conditions, it does not necessarily mean that the ParA_F_ oscillation inherently hinges on nucleoid splitting. Rather, in more physiologically relevant condition as in our experiments, ParA_F_ oscillations occur independently of nucleoid division and PC movement.

Our observations precipitate many open questions, pointing to the heart of the physical mechanism that underlies the ParAB*S*-mediated genome partition. For instance, what drives the ParA_F_ oscillation that is not tightly coupled to PC movements? Moreover, this ParA_F_ pole-to-pole oscillation results in a large variation in the ParA_F_ intracellular concentration inherited by the daughter cells upon cell division (15). While the PC localizes ParA_F_ nearby to buffer against this large cell-to-cell variation in ParA_F_ level, underlying robust PC partition (15), it is not understood why the ParA_F_ needs to oscillate after all. Particularly, given that ParA and ParB gene expression are autoregulated (29, 30), why doesn’t the cell prevent the ParA_F_ oscillation and quench the large cell-to-cell variation in ParA_F_ intracellular concentration? Wouldn’t the latter strategy better ensure the fidelity of PC partition? We next combined theoretical modeling with experiment to address these pressing questions.

### Model development

We aimed to construct a minimal model to recapitulate the essence of the ParA_F_ oscillation observed in our experiments (**Fig. 1**). This model integrates the essential interactions between ParA_F_ and ParB_F_ constrained by the existing biochemical studies: *i.e.*, ParA_F_ promotes the nucleoid-binding for itself via positive feedback; and the nucleoid-bound ParA_F_ promotes the nucleoid-binding of ParB_F_, which in turn triggers ParA_F_ dissociation from the nucleoid, constituting negative feedback (**Fig. 2A**). As shown below, this intertwined feedback mechanism is sufficient to explain the observed ParA_F_ oscillation.

**Fig. 2.**
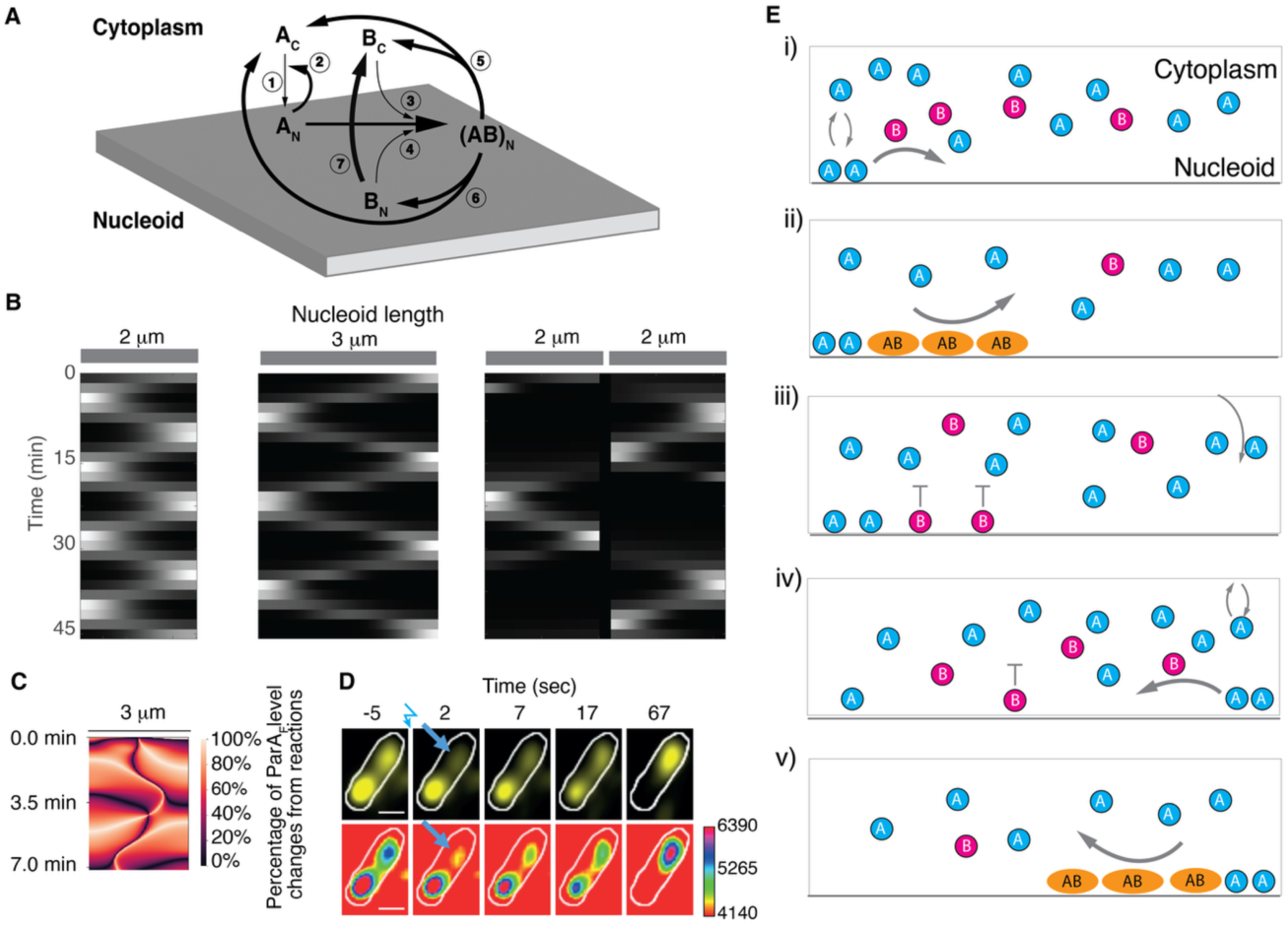
Theoretical model captures the essence of observed ParA_F_ oscillation. A) Model schematics of ParA_F_/ParB_F_ biochemical reactions. Hereby, ParA_F_ binding to nucleoid is self-reinforcing as positive feedback (Reactions #1 and 2). In contrast, the nucleoid-bound ParA_F_ recruits cytosolic ParB_F_ or nucleoid-bound ParB_F_ to form ParA_F_–ParB_F_ complex on nucleoid (Reactions #3 and 4). The ParA_F_–ParB_F_ complex rapidly turns over via two pathways. The first pathway is that nucleoid-bound ParA_F_–ParB_F_ complex directly dissociates into the cytosolic ParA_F_ and ParB_F_ (Reaction #5). The second pathway is that nucleoid-bound ParA_F_–ParB_F_ complex dissociates into the cytosolic ParA_F_ and the nucleoid-bound ParB_F_ (Reaction #6). Further, the nucleoid-bound ParB_F_ will dissociate into cytosol very rapidly (Reaction #7). The reactions #3-7 together constitute negative feedback, in which the nucleoid-bound ParA_F_ essentially promotes its own dissociation from the nucleoid. The ParA_F_ oscillation hinges on the above intertwined feedback. The model does not explicitly depict the PC nor its movement, since they are not tightly coupled to the ParA_F_ oscillation as evidenced in our experiments. Instead, the model assumes that the PC controls the level of the available ParB_F_ molecules for cytosolic diffusion and nucleoid-binding away from the PC, because the PC is known to sequester the majority of ParB_F_ inside the cell. B) Kymographs of ParA_F_ oscillation from model results on three different nucleoid lengths. C) ParA_F_ oscillation is a phase wave. The chemical reactions constitute > 98% of the total changes in the local nucleoid-bound ParA_F_ concentration. D) A majority of ParA_F_ redistributes on the other cell-half during oscillation on live cell. Time-lapse microscopy images from a FRAP experiment in which the cell-half exhibiting the lowest ParA_F_ fluorescence was photobleached at time 0. The same cell is displayed in the yellow channel (*top*) or in heat map (*bottom*). The blue zigzag symbol and the blue arrow indicate the timing and location of photobleaching, respectively. Time is indicated in seconds. Scale: 1 µm. E) Schematic elucidating that the spatial separation between ParA_F_ and ParB_F_ underlies the see-saw oscillation.

In contrast to the previous model that relies on plasmid interactions or physical motion to facilitate ParA_F_ oscillation, our model does not explicitly describe the F-plasmids, because our data suggest no direct correlation between plasmid mobility and ParA_F_ oscillation (**Fig. 1F**). Given that over 90% of ParB_F_ is condensed around the *parS*_F_ centromere site inside *E. coli* cells (43, 44), the model implicitly depicts the effect of F-plasmids as controlling the free ParB_F_ molecules that can either diffuse in the cytosol or interact with the nucleoid-bound ParA_F_.

Mathematically, we describe the dynamics of ParA_F_–ParB_F_ interactions with a set of reaction-diffusion evolving over two juxtaposed 2D simulation domains representing the cytosol and nucleoid, respectively. While freely diffusing within the each of the two simulation domains, the ParA_F_ and ParB_F_ molecules exchange between cytosol and nucleoid when the chemical reactions take place. We ignore the plasmid and plasmid-bound ParB_F_, only considering concentrations of free ParA_F_ and ParB_F_, as well as independent ParA_F_-ParB_F_ complexes on the nucleoid. The model is composed of three key elements:

1) ParA_F_ accumulates on the nucleoid in clusters (14) via a positive feedback loop. Cytosolic ParA_F_ independently, nonspecifically binds to the nucleoid, where consequently it recruits additional ParA_F_ from the cytosol via cooperative binding (35, 45), leading to local accumulation.
2) Accumulating ParA_F_ is depleted from the nucleoid by recruiting and interacting with ParB_F_. This negative feedback loop consists of ParA_F_-ParB_F_ complex formation and dissociation. Once nucleoid-bound, the ParA_F_ recruits diffusive ParB_F_ to form a ParA_F_-ParB_F_ complex that, upon dissociation, will release ParA_F_ from the nucleoid into the cytosol (5). The dissociated return to diffusive ParB where it can be recruited by a different nucleoid-bound ParA_F_ to form another ParA_F_-ParB_F_ complex again (5). Additionally, ParB_F_ has an independent dissociation rate from the nucleoid (6).
3) ParA_F_, ParB_F_, and ParA_F_-ParB_F_ complex diffuse much more slowly on nucleoid than in cytosol (14, 44).

We would like to stress from the outset that to discern the fundamental physical picture of the ParA_F_ oscillation wave, the model has several simplifications of the detailed ParABS biochemistry. And importantly, these simplications do not notably alter the essential model results, based on the quantitative considerations from existing experimental measurements. First, the ParA_F_ in the model refers to the ParA_F_ dimer and, hence, does not describe the ParA_F_ monomer to dimer transition, because such a transition is much faster than the temporal resolution of the model (5). This model simplification is consistent with the prior studies showing that unlike chromosomally-encoded ParA, ParA_F_ is dimeric in solution at physiological concentrations (46, 47). Second, our model does not consider the ParA_F_-independent ParB_F_-nucleoid binding. This is because, in the absence of ParA the ParB-nucleoid binding requires a much higher cytosolic ParB_F_ concentration than that *in vivo*, and the nucleoid-bound ParB displays a very high off rate (48). Third, the model ignores the ParB_F_-independent ParA_F_ dissociation from the nucleoid, because it is much slower than the ParB_F_-dependent counterpart (*i.e.*, 1.9 min^-1^ *vs.* 8 min^-1^, see the Fig. 1F in (6)). Indeed, our model calculation shows that the essence of ParA oscillation wave persists, as long as the ParA_F_-independent ParB_F_-nucleoid binding is slower than the ParA_F_-dependent counterpart and the ParB_F_-independent ParA_F_ dissociation has a minor contribution to the overall ParA-nucleoid dissociation event. Lastly, the model does not explicitly depict the observed time-delay, by which a ParA molecule regains the nsDNA-binding capacity once it dissociates from the nucleoid (6). Instead of tracking the individual ParA at a single-molecule level, the model captures the essence of the time-delay at a collective level: *i.e.*, the local rate of cytosolic ParA-nucleoid binding is very slow when there is no nucleoid-bound ParA at this location; consequently and conversely, once the ParA dissociates from the nucleoid, it will take a while for the cytosolic ParA to bind the nucleoid at this location.

The coupled 2D reaction-diffusion equations of the model are as follows:

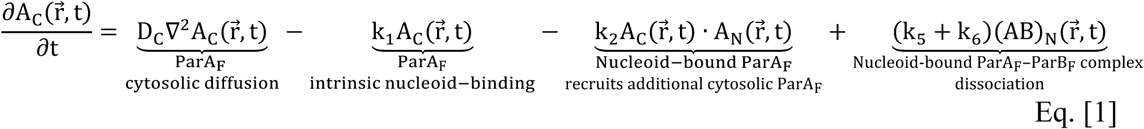

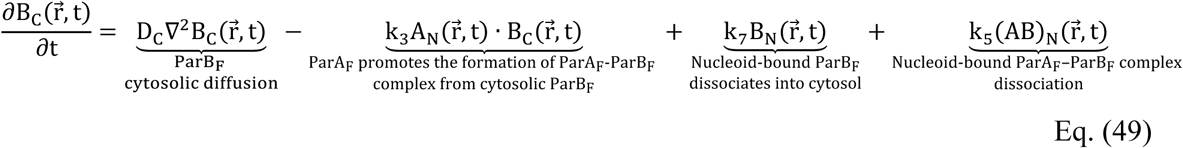

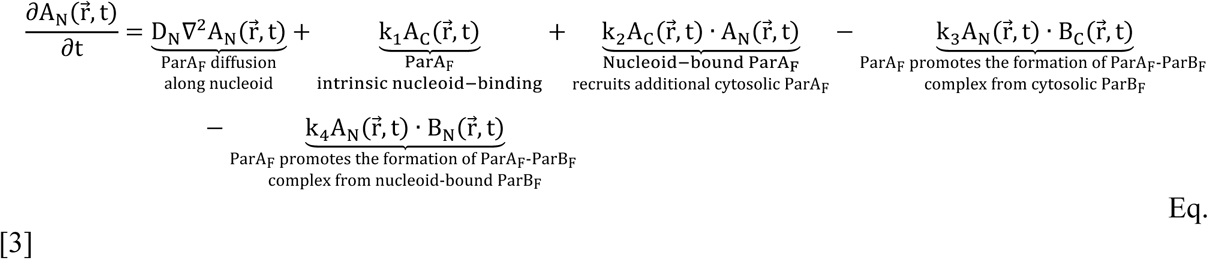

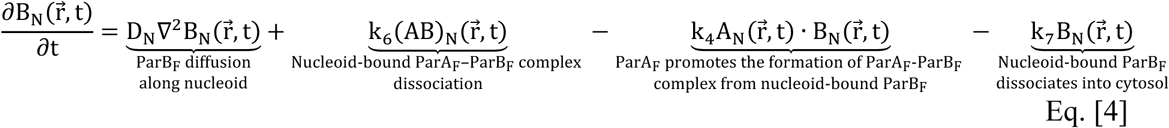

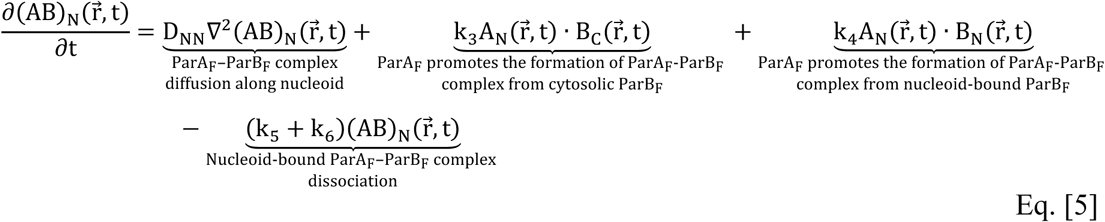

Here, A_C_(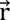, t) and B_C_(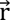, t) are the cytosolic concentrations of ParA_F_ and ParB_F_ at the position, 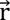 and the time, t, respectively. Likewise, A_8_(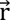, t) and B_8_(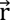, t) are the corresponding nucleoid-bound concentrations, and (AB)_8_(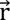, t) represents the concentration of ParA_F_-ParB_F_ complex on the nucleoid. The reaction and diffusion processes are briefly depicted underneath each term in the Eqs. [1-5]. All the model parameters are derived from the experimental measurements whenever possible and listed in the **Table 1**. If not otherwise mentioned, the ParA_F_ and ParB_F_ concentrations in the simulation were kept fixed as the nucleoid elongates and splits. Our nominal case of the model was based on our experimental measurements (43, 50, 51): the fixed concentrations correspond to 1250 ParA_F_ molecules and 225 ParB_F_ molecules on a 3 μm-by-1 μm nucleoid, respectively. The initial condition was chosen so that the symmetry was already broken, *i.e.*, all the ParA_F_ and ParB_F_ were in the left half of the nucleoid. We use an elementary finite-difference Euler scheme to calculate the concentration differences at each timestep. For computational efficiency, we calculate the diffusive component at each timestep using a Crank-Nicholson method to separately calculate and combine the diffusive contribution at the x and y axis. To capture the observed essence that the ParA_F_ oscillation evolves as the nucleoid elongates and splits during cell division, we adapted the model to the nucleoids at the different lengths.

**Table 1.** Model Parameter.

| Parameters | Physical meaning | Measured values | Nominal values |
| --- | --- | --- | --- |
| $D_c$ | Diffusion constant of ParA<sub>F</sub> and ParB<sub>F</sub> in cytosol | $\sim 1 \mu\text{m}^2/\text{s}$ (44, 58) | $0.5 \mu\text{m}^2/\text{s}$ (\$) |
| $D_n$ | Diffusion constant of ParA<sub>F</sub> and ParB<sub>F</sub> on nucleoid | $\sim 10^{-4} - 10^{-2} \mu\text{m}^2/\text{s}$ (6, 58) | $2.5 \times 10^{-3} \mu\text{m}^2/\text{s}$ (\$) |
| $D_{nn}$ | Diffusion constant of ParA<sub>F</sub>-ParB<sub>F</sub> complex on nucleoid | half of $D_n$ | $1.3 \times 10^{-3} \mu\text{m}^2/\text{s}$ (\$) |
| $k_1$ | Intrinsic nucleoid-binding rate of ParA<sub>F</sub> | $\sim 10^{-4} - 10^{-2} / \text{s}$ (3, 5, 6, 11, 45) | $7.1 \times 10^{-4} / \text{s}$ (&) |
| $k_2$ | ParA<sub>F</sub> nucleoid-binding rate in the positive feedback | Estimated | $8.5 \times 10^{-4} \mu\text{m}^2/\text{s}$ (*) |
| $k_3$ | Formation rate of ParA<sub>F</sub>-ParB<sub>F</sub> complex from nucleoid-bound ParA<sub>F</sub> and cytosolic ParB<sub>F</sub> | Estimated | $1.1 \times 10^{-2} \mu\text{m}^2/\text{s}$ (*) |
| $k_4$ | Formation rate of ParA<sub>F</sub>-ParB<sub>F</sub> complex from nucleoid-bound ParA<sub>F</sub> and ParB<sub>F</sub> | Estimated | $2.3 \times 10^{-2} \mu\text{m}^2/\text{s}$ (*) |
| $k_5$ | Rate of ParA<sub>F</sub>-ParB<sub>F</sub> complex dissociating into the cytosolic ParA<sub>F</sub> and the nucleoid-bound ParB<sub>F</sub> | $\sim 0.05 - 1 / \text{s}$ (6) | 0.07/s |
| $k_6$ | Rate of ParA<sub>F</sub>-ParB<sub>F</sub> complex dissociating into the cytosolic ParA<sub>F</sub> and ParB<sub>F</sub> | $\sim 0.05 - 1 / \text{s}$ (6) | 0.07/s |
| $k_7$ | Rate of nucleoid-bound ParB<sub>F</sub> dissociating into cytosol | $\sim 0.1 - 1 / \text{s}$ (48, 59) | 0.95/s |

Note (\$): In the model, the diffusion constant of ParA<sub>F</sub> and ParB<sub>F</sub> molecules in cytosol, $D_c$ corresponds to the experimentally measured diffusion constant of the protein mutants that do not bind to the nucleoid, diffuse freely in cytosol, and are consistent with the expected diffusion constant of an inert particle with the similar size ( $\sim 1 \mu\text{m}^2/\text{s}$ ) (58). In contrast, the diffusion constant of the nucleoid-bound ParA<sub>F</sub> and ParB<sub>F</sub> molecules should correspond to the measured one in WT by FRAP and other microscopies (6, 58), which is in the range of $\sim 0.01 - 0.1 \mu\text{m}^2/\text{s}$ . Furthermore, the ParA<sub>F</sub>-ParB<sub>F</sub> complex is ∼ twice in size of either ParA<sub>F</sub> or ParB<sub>F</sub>; hence, its diffusion constant on nucleoid is assumed to be half of that for nucleoid-ParA<sub>F</sub> or ParB<sub>F</sub>.
Note (&): The experiments measured the apparent nucleoid-binding rate of ParA<sub>F</sub> to be $\sim 0.01 - 0.1 / \text{s}$ (3, 5, 6, 11, 45), which includes both the intrinsic rate, $k_1$ and that from the positive feedback. Given that the positive feedback constitutes the majority of the ParA<sub>F</sub> nucleoid-binding (5), the model chooses the value of $k_1$ to be order-of-magnitude smaller than the measured apparent rate of ParA<sub>F</sub> nucleoid-binding.
Note (\*): These chemical reaction rates are associated with the second-order reactions on a 2D simulation domain, where the protein concentration corresponds to the surface density (*i.e.*, the number of protein per area).

### Theoretical model captures the observed ParA_F_ oscillation as a see-saw phase wave

**Fig. 2B** shows the typical model results. First, the ParA_F_ oscillates along the coherent nucleoid of different lengths (e.g., 2 μm and 3 μm in the first two panels of **Fig. 2B**). Second, such an oscillation persists when the nucleoid splits (4 μm, the third panel of **Fig. 2B**). Third and importantly, the emergence of the ParA_F_ oscillation in the model does not hinge on the nucleoid segregation nor the PC movement. These features are consistent with our observations (**Fig. 1**).

Our model further predicts that the ParA_F_ oscillation does not reflect the diffusional drift of ParA_F_ along the nucleoid surface with a sustained wavefront. Rather, it displays a see-saw pattern with the decreased ParA_F_ intensity at the middle ground, as the ParA_F_ rapidly switches between the two halves of the nucleoid (**Fig. 2B**). Note, the ParA_F_ intensity significantly decreases at the middle ground of the nucleoid during the ParA_F_ oscillation regardless of whether the nucleoid splits or not, although the intensity drop is deeper when the nucleoid splits (**Fig. 2B**). Importantly, the model predicts that the changes in their nucleoid-bound concentrations mainly reflect the recruitment from cytosol, rather than the diffusional drift of the proteins along the nucleoid (**Fig. 2C**). In other words, the ParA_F_ oscillation is a phase wave.

These predicted features are consistent with our kymograph measurements, which reveals the very dim ParA_F_ intensity at the mid-nucleoid as the ParA_F_ oscillate from pole to pole (**Fig. 1C**). This indicates that during oscillation the majority of ParA_F_ molecules does not traverse across the middle ground along the nucleoid surface. Further, this see-saw pattern is supported by our FRAP experiments (**Fig. 2D**): During the ParA_F_ oscillation, the rapidly-photobleached low-ParA_F_ nucleoid half “attracted” most of the ParA_F_ molecules from the cytosol above the other nucleoid half, while the ParA_F_ intensity at the mid-ground remained low. Combining the data in **Figs. 1C** and **2D**, it suggests that during oscillation, the majority of nucleoid-bound ParA_F_ redistribution along the nucleoid surface is mediated through the cytosolic pool.

The theory and experimental results in **Figs. 1C** and **2B-D** led us to the following physical picture of ParA_F_ oscillation (**Fig. 2E**): the see-saw pattern stems from the ParA_F_ intensities on the two nucleoid halves act as two spheres of power that mainly communicate through the ParA_F_ diffusion in the cytosol, rather than the diffusional drift across the mid-ground of nucleoid. Specifically, as ParA_F_ rapidly binds onto the left half of the nucleoid via the positive feedback, the accumulation of the nucleoid-bound ParA_F_ promotes the ParB_F_ binding onto the nucleoid. When these nucleoid-bound ParB_F_ starts to “kick” off the ParA_F_ locally, the dissociated ParA_F_ can diffuse rapidly in the cytosol both in the forward and backward directions. At this stage, plenty of the ParA_F_ still remains on the left half of the nucleoid and retains the ParB_F_ molecules, which prevent the dissociated ParA_F_ from rebinding. In contrast, the right half of the nucleoid is largely devoid of ParB_F_ molecules and, hence, favors the ParA_F_ nucleoid-binding from the cytosol. Given that the closed boundary condition prevents the cytosolic proteins from diffusing away, more ParA_F_ molecules bind to the right half of the nucleoid. Such a spatial separation between ParA_F_ and ParB_F_ allows the ParA_F_ positive feedback to take over on the right, which further attracts more cytosolic ParA_F_ from the left. Eventually, the culmination of ParA_F_ binding onto the right half of the nucleoid tips the balance and depletes the nucleoid-bound ParA_F_ on the left. This causes the rapid switch of the nucleoid-bound ParA_F_ concentration from the left to the right halves of the nucleoid, which in turn repeats the above episodes, driving the ParA_F_ pole-to-pole oscillation.

We next asked why the ParA_F_ intensity decreases at the middle ground of the nucleoid during the ParA_F_ oscillation? Our model suggests that the fundamental reason lies in the residual concentration of nucleoid-bound ParB_F_ at the mid-nucleoid, shaped by the reaction-diffusion process with the closed boundary at the cell poles (**Fig. 2E**). Consider the stage that the majority of ParA_F_ and ParB_F_ binds to the left half of the nucleoid. Albeit slowly, the nucleoid-bound ParB_F_ can diffuse to the middle ground and prevents the local nucleoid-binding of cytosolic ParA_F_. In contrast, the right half of the nucleoid has a very low level of ParB_F_ and, hence, “attracts” the ParA_F_ via the cytosolic diffusion. Importantly, the closed boundary at the nucleoid end prevents the cytosolic ParA_F_ from diffusing away. This way, the ParA_F_ keeps binding to the right half of the nucleoid via the positive feedback, while the ParB_F_ negative feedback takes over at the mid-nucleoid. As a natural consequence of the positive feedback, the high-ParA_F_ nucleoid region will attract more cytosolic ParA_F_ at the expense of the low-ParA_F_ nucleoid region, when the overall ParA_F_ level is limited but still supports the oscillation (as in our nominal model parameter set (**Fig. 2**)). This further increases the nucleoid-bound ParA_F_ level at the right half of the nucleoid, relative to that at mid-nucleoid. Therefore, during the ParA_F_ oscillation the concentration of the nucleoid-bound ParA_F_ is lower at the mid-nucleoid than the peak concentration on either half of the nucleoid. In other word, the see-saw pattern arises from the combination of three key factors: the close boundary condition, the ParA_F_ positive feedback, and the ParB_F_ negative feedback.

So far, our work suggests that the ParA_F_ oscillation along the nucleoid reflects a phase wave, which switches from pole to pole with little ParA_F_ physically traversing along the nucleoid through the middle ground . And the model-experiment agreement provides a solid foundation for further model predictions. Our model indicates that the ParA_F_ oscillation intrinsically hinges on the geometry of the nucleoid and intertwined feedback between ParA_F_ and ParB_F_ in nucleoid-binding. Combining experiments and modeling, we next tested these predictions to determine the physical mechanisms of the ParA_F_ oscillation.

### ParA_F_ oscillation critically depends on nucleoid dimension

To examine how nucleoid geometry influences the ParA_F_ see-saw oscillation, we first exploited a RodZ-deficient mutant to experimentally alter cell shape and hence nucleoid morphology (**Fig. 3A**). In this mutant, the normally rod-shaped E. coli cells become spherical, leading to a round nucleoid. **Fig. 3B** shows that in rounded cells with a low length/width ratio (aka aspect ratio), the characteristic ParA_F_ oscillation is absent. Interestingly, as these spherical cells elongate during cell division, the ParA_F_ oscillation re-emerges transiently once the aspect ratio of the cell becomes greater than 1.6 (N=15), before disappearing again upon cell division, when the two rounded daughter cells are formed. We next leveraged our model to understand how the ParA_F_ oscillation depends on the geometry of the nucleoid. To meaningfully engage with our RodZ mutant experiment, we first altered the aspect ratio of the nucleoid in the model by fixing the width at 1 μm and increasing the length, while keeping the concentration of ParA_F_ and ParB_F_ fixed. Consistent with experiments, the model results show that the ParA_F_ oscillation emerges only when the aspect ratio of the nucleoid is above > 1.3, *i.e.*, the nucleoid is sufficiently elongated (**Fig. 3C**).

**Fig. 3.**
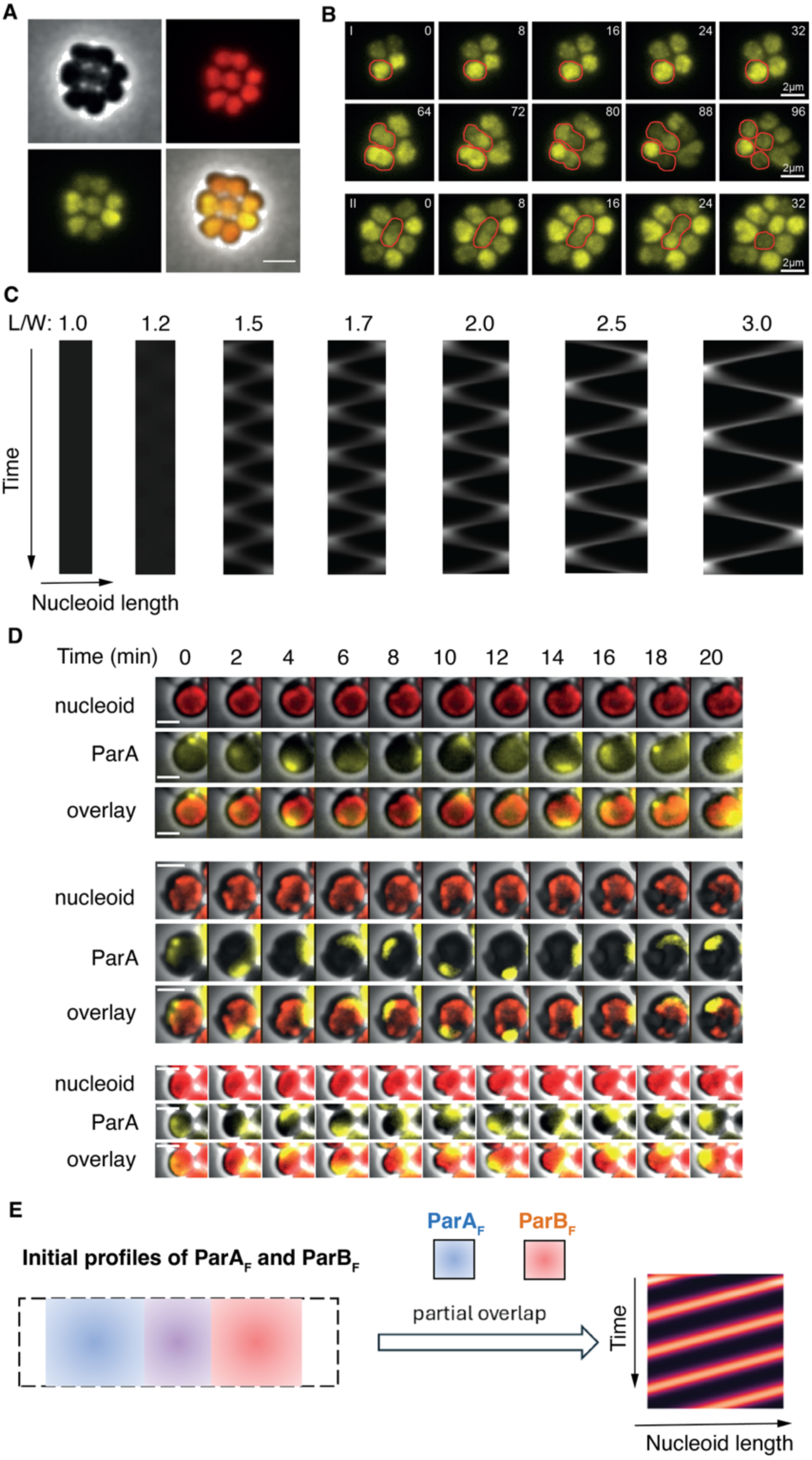
Nucleoid geometry controls the ParA_F_ oscillation. A) Images of RodZ mutant cells in bright field, red channel (nucleoid/Hu-mCherry), yellow channel (ParA-mVenus) and combined. B) Time lapse microscopy images (I-II) showing rounded cells that start ParA_F_ oscillations when the aspect ratio (length/width) is above 1.6 (N=15) and resume oscillation after cell division gave rise to two rounded cells. Time is indicated in min. C) Model results capture the dependence of ParA_F_ oscillation on nucleoid shape. D) ParA_F_ pole-to-pole oscillation changes to circulating wave when the nucleoid becomes a donut-shape upon A22 treatment. Time-lapse imaging of rounded cells after treatment with A22. Images were taken every 30 sec over 30 min. in bright field, red, and yellow channels to observe the cell contour, the Hu-mCherry labelled nucleoid, and ParA_F_-mVenus, respectively. For simplicity, images are displayed every 2 min for 20 min. In the three typical examples displayed, ParA_F_ circulates endless over the rounded nucleoid. Scale bar: 2 µm. E) Model results with periodic boundary condition recapitulate the observed circulating wave. *Left*: Initial condition that ParA_F_ and ParB_F_ concentration fields partially overlap. *Right*: Kymograph from model simulation showing that ParA_F_ displays circulating wave with the periodic boundary condition.

This finding precipitates a further question: What defines this threshold nucleoid length? As elaborated above, the necessary condition of the ParA_F_ see-saw oscillation is the spatial separation between the nucleoid-bound ParA_F_ and ParB_F_ (**Fig. 2E**). This spatial separation (L_S_) arises from the lag time (τ) between the initiation timing of the local ParA_F_ positive feedback and that of ParB_F_ negative feedback. Heuristically, L_S_ should scale with the nucleoid length and the boundary condition, and τ should scale with the oscillation period. Importantly, this lag time defines a characteristic length scale of the system, 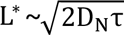, which is the diffusion distance that the nucleoid-bound ParA_F_ and ParB_F_ traverse along the length of the nucleoid ( D_N_ is their diffusion constant). If L^*^is comparable or longer than the L_S_ , then the separation between the nucleoid-bound ParA_F_ and ParB_F_ will be homogenized by the protein diffusions along the nucleoid. Hence, the oscillation will be disrupted.

### A donut-shaped nucleoid supports a circulating ParA_F_ wave

The above mechanistic insight leads to an interesting question: How does ParA_F_ oscillation respond if the nucleoid keeps elongating without divisions? Although it is experimentally challenging to prevent the elongating nucleoid from splitting in experiments, we exploited A22 treatment to perturb cell morphology. Fluorescent imaging revealed that some nucleoids adopt a donut-shape upon A22 treatment (**Fig. 3D**). We examined the PC positioning on these donut-shaped nucleoids (**Fig. S3**) and found that they remained regularly spaced relative to one another. Such spacing is consistent with active partitioning, indicating that the partition machinery remains functional despite the altered geometry. Surprisingly, instead of the typical pole-to-pole oscillation, ParA_F_ displayed a circulating wave that continuously propagated along the perimeter of the donut-shaped nucleoid (**Fig. 3D**). This donut-shaped nucleoid essentially translates into a periodic boundary condition in the model, thus representing an infinite long nucleoid.

This unexpected observation provides a testing ground of our proposed physical mechanism. We therefore asked whether our model with the lengthwise periodic boundary condition could capture the essence of the observed circulating wave? The key lies in the spatial separation between the ParA_F_ and ParB_F_. In our model, a partial overlap in their initial distributions of ParA_F_ and ParB_F_ can readily drive a circulating wave (**Fig. 3E**). Consider that ParA_F_ initially accumulates more toward the left half of the nucleoid, for instance. This distribution already breaks the symmetry, promotes more local ParA_F_ nucleoid-binding, and sets the wave propagation to the left. Since there is no closed boundary in the circular case, the cytosolic ParA_F_ can diffuse freely without being stalled. This is in a stark contrast to our WT case, in which ParA_F_ builds up the high-density near the nucleoid end and the low-density at the mid-nucleoid, resulting in the see-saw patterns (**Fig. 1 and 2**). Instead, the deposition of the cytosolic ParA_F_ on the circular nucleoid keep propagating with a constant density at the wavefront. Hereby, the ParA_F_-mediated positive feedback dominates at the propagating wave front by self-promoting the nucleoid-binding, whereas the ParB_F_-mediated negative feedback dissociates the ParA_F_ at the trailing tail of the wave. It is such a spatial separation that sustains the wavefront of a constant ParA_F_ density circulating around the circumference of the nucleoid (**Fig. 3E**).

Combining the experiment and theoretical modeling, our finding establishes that the nucleoid geometry critically modulates ParA_F_ oscillation. According to our model, while the nucleoid length and boundary conditions define the geometry context, it is the feedback between ParA_F_ and ParB_F_ nucleoid-binding that underlies the molecular mechanism for the ParA_F_ oscillation. We next examined this feedback and its potential functions.

### ParB_F_-mediated negative feedback controls ParA_F_ oscillation

The model predicts that due to the intertwined feedback between ParA_F_ and ParB_F_, the average level of the nucleoid-bound ParA_F_ depends on the ParA_F_ to ParB_F_ ratio in a highly cooperative manner (**Fig. 4A**). When ParA_F_ to ParB_F_ ratio is very high, the ParA_F_ positive feedback dominates and most of the intracellular ParA_F_ molecules binds to the nucleoid. As the ParA_F_ to ParB_F_ ratio decreases, the ParB_F_-mediated negative feedback starts to manifest its effect and consequently, the average level of the nucleoid-bound ParA_F_ starts to decrease. Crucially, when the ParA_F_ to ParB_F_ ratio decreases a merely 2 folds (from 4 to 2), the average level of the nucleoid-bound ParA_F_ sharply decreases more than 100 folds. These model results – that hinge on the ParB_F_-mediated negative feedback – provide a mechanistic underpinning for the corresponding titration curve observed by the *in vitro* experiments (6). Intriguingly, the ParA_F_ oscillation exists around the transition regime of the sigmodal curve (**Fig. 4A**), where the ParB_F_-mediated negative feedback becomes notable but not predominant over the ParA_F_ positive feedback.

**Fig. 4.**
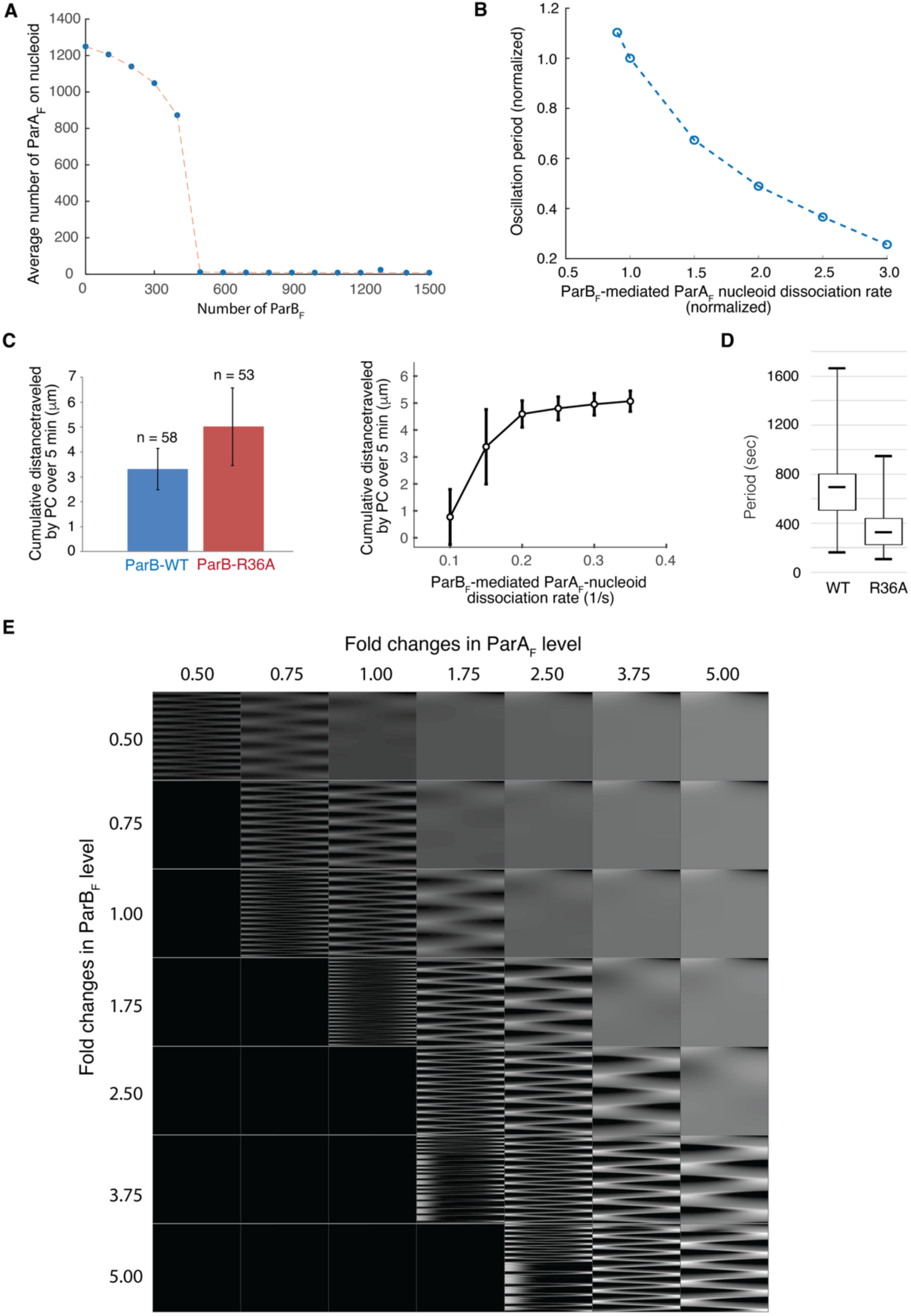
Negative feedback between ParA_F_ and ParB_F_ controls ParA_F_ oscillation and fidelity of F-plasmid partition. A) The model results recapitulate the observation that ParB_F_ controls the average level of nucleoid-bound ParA_F_. In this model calculation, only the total ParB_F_ concentration was varied with all the other parameters fixed as those in the nominal case (**Fig. 2B**); in particular, the nucleoid length was 3 μm and the total concentration of ParA_F_ was kept so that there were ∼ 2500 ParA_F_ molecules in the cell with 3 μm-nucleoid. B) The model predicts that ParB_F_-driven ParA_F_ dissociation rate from nucleoid controls the period of ParA_F_ oscillation. C) ParA_F_ nucleoid dissociation rate increases in ParB_F_-R36A mutant. *Left*: Experimental data of cumulative distance in WT and ParB_F_-R36A mutant. *Right*: Model result on the dependence of cumulative distance on koff with the agent-based stochastic simulation of PC partition. We fit the model (*right*) with the experimental data (*left*) to infer the increase in the ParB_F_-mediated dissociation rate of ParA_F_ from DNA. D) Period of ParA_F_ oscillation shortens in ParB_F_-R36A mutant as compared to WT. E) Model phase diagram calculation predicts that ParA_F_ oscillation requires a proper ratio between ParA_F_ and ParB_F_. In the model calculation, we varied the ParA_F_ and ParB_F_ concentrations, while keeping all the other parameters, including the nucleoid length at 3 µm and implementation of the closed boundary condition. In the phase diagram, we plotted the kymograph to characterize the spatial-temporal pattern of nucleoid-bound ParA_F_ for each pair of ParA_F_ and ParB_F_ concentrations.

In addition to the intracellular level of ParB_F_ molecules, the ParB_F_-mediated dissociation rate of ParA_F_ from the nucleoid is another key parameter controlling the associated negative feedback. Following this line of reasoning, the model predicts that increasing the ParB_F_-mediated dissociation rate of ParA_F_ from the nucleoid shortens the period of ParA_F_ oscillation within the oscillation regime (**Fig. 4B**).

To experimentally test this prediction, we resorted to the ParB_F_-R36A mutant that is proficient in ParA-ParB interactions but defective in ParB_F_ stimulated ParA ATPase activity (52) and, hence, expected to influence the ParA_F_-nucleoid dissociation rate. Indeed, the partition complexes (PCs) became much more mobile than the WT (**Fig. 4C**, left panel) – the cumulative displacement (CD) of the PCs is much larger in ParB_F_-R36A mutant than in WT (13). To quantify how the ParA_F_-nucleoid dissociation rate changes in the ParB_F_-R36A mutant, we exploited our previous Brownian ratchet model of ParABS-mediated PC partition (9, 10, 15) and calculate how the koff controls the CD of the PCs as a titration curve (**Fig. 4C**, right panel). To infer the changes in the koff, we then fitted the experimentally measured CD to this model result. Our fitting result shows that the koff increases by ∼ 100 % in ParB_F_-R36A mutant. With the ∼ 100% increase of koff, the model further predicts that the period of ParA_F_ oscillation will shorten by > 2 folds (**Fig. 4B**). Indeed, quantifying our experimental data of the ParA_F_ oscillation shows that the oscillation period (13), on average, shortens by ∼ 2 folds, quantitatively supporting our model prediction.

To have a more global view of the ParA_F_ oscillation, we next calculated our model phase diagram to determine how it depends on the intracellular concentrations of ParA_F_ and ParB_F_ molecules (**Fig. 4E**). The phase diagram predicts that ParA_F_ oscillation requires a proper ratio between ParA_F_ and ParB_F_ (**Fig. 4E**): When the ParA_F_ level is too high or too low, it disrupts ParA_F_ oscillation. Specifically, at a high ratio of ParA_F_ to ParB_F_, the nucleoid is saturated with a uniform coat of ParA_F_, undergoing no oscillation. Here, ParA_F_ positive feedback outcompetes ParB_F_’s ability to trigger dissociation of ParA_F_ from the nucleoid. However, when the ratio of ParA_F_ to ParB_F_ is low, an excess of ParB_F_ will quench the cooperative properties of ParA_F_ binding to the nucleoid, triggering nucleoid dissociation before additional ParA_F_ are recruited, leading to an effectively vacant nucleoid, save for transient binding/unbinding. The oscillating region is within a critical range of ParA_F_:ParB_F_ ratio, visualized along the main diagonal of the phase diagram.

To experimentally test this prediction, we increased the intracellular ParA_F_ level while maintaining ParB_F_ levels relatively constant. For this, the *E. coli* cells carrying the mini-F plasmid were transformed with an additional plasmid expressing *parA*_F_ under the control of an arabinose-inducible promoter. We quantified intracellular ParA_F_ and ParB_F_ levels at various concentrations of inducer by Western blot analyses (**Fig. S4A-B**) and measured alongside both the oscillation periods of ParA_F_ and the partition efficiency (Loss rate) of plasmid F (**Table 2** and **Fig. S4C**). As predicted (**Fig. 4E**), a moderate increase in the intracellular ParA_F_ level (ParA_F_/ParB_F_ ratio ∼1.7) slowed down the oscillation, whereas a large excess (> 14) abolished it. While a 2-fold increase in the ParA/ParB ratio did not affect plasmid F stability, higher ratios (∼14 and ∼22) significantly reduced or abolished partition fidelity, respectively. These results indicate that the ParA/ParB ratio is essential for accurate partitioning and that loss of oscillation correlates with partition defects. Together, these findings support a model in which a negative feedback loop between ParA_F_ and ParB_F_ controls the dynamics of ParA_F_ oscillation.

**Table 2.** Plasmid loss rates under varying levels of partition proteins.

| Plasmids | Ara (%) | Loss rate (%) | Relative ParA | Relative ParB | ParA/ParB ratio | ParA oscillation period |
| --- | --- | --- | --- | --- | --- | --- |
| pDAG114 | - | < 0.01 | nd | nd | na | na |
| pDAG115 | - | 5.75 | nd | nd | na | na |
| pJYB243 | 0 | <0.01* | 1 | 1 | 1 | 13 min |
| pJYB243+pJYB353 | 0 | <0.01 | 0.68 | 0.71 | 0.96 | 13 min |
| pJYB243+ pJYB353 | 10<sup>-4</sup> | <0.01 | 0.85 | 0.48 | 1.77 | 17 min |
| pJYB243+ pJYB353 | 10<sup>-3</sup> | 0.65 | 4.89 | 0.34 | 14.4 | No oscillation |
| pJYB243+ pJYB353 | 3×10<sup>-3</sup> | 1.10 | 7.78 | 0.45 | 17.3 | No oscillation |
| pJYB243+ pJYB353 | 10<sup>-2</sup> | 4.90 | 11.1 | 0.50 | 22.2 | No oscillation |

*nd*, not determined; *na*, not applicable

### ParB_F_-mediated negative feedback controls PC partition fidelity

Lastly, we asked what functional role ParA_F_ oscillation plays in plasmid partitioning. Our data show that overexpression of ParA_F_, which abolishes oscillation, leads to a drastic increase in plasmid loss rate (**Table 2**). In contrast, partition accuracy remained largely unchanged under conditions that preserve the ParA_F_ oscillation, including (i) a moderate ParA_F_ overexpression that slows oscillation (**Table 2**) and (ii) with the ParB_F_-R36A mutant, which speeds up oscillation (13). These findings suggest that ParA_F_ oscillation may relate to the fidelity of PC partition.

According to our ParA concentration gradient-based Brownian ratchet model of PC partition (7, 9, 10, 14, 15, 17), when the nucleoid-bound ParA_F_ level is too high, the replicated PCs segregate with a shortened distance. Given that the timing of PC replication is not tightly coupled to the cell division (53) and that PC is constantly on the move, this shortened segregation distance renders a higher chance that the two PCs end up in the same half of the nucleoid during cell division. Conversely, when the nucleoid-bound ParA_F_ level is too low, the PC simply undergoes diffusion with about 50% percentage of chance that a daughter cell inherits both PCs. Either way, the nucleoid-bound ParA_F_ level needs to be properly controlled to ensure the partition fidelity (7, 9, 10, 14, 15, 17). From this perspective, our work shows that the ParB_F_-mediated negative feedback provides a control over the nucleoid-bound ParA_F_ level (**Fig. 4**), and the ParA_F_ oscillation may be the readout of this elaborate control. Further, this is a robust control in that the ParA_F_ oscillation remains insensitive against a variety of variations in growth conditions (**Fig.1**).

Taken together, our data is consistent with the view that the ParB_F_-mediated negative feedback is robust in supporting the PC partition fidelity by buffering against parameter variations, and that the ParA_F_ oscillation wave may be reminiscent of this robust molecular mechanism.

## Discussion

Combining experiment and theoretical modeling, we present evidence that the ParA is excitable during cell division. It oscillates from pole to pole along the nucleoid length in a see-saw pattern. Hereby, instead of representing the real material propagation in the direction of the wave, the apparent wave mainly reflects the cytosolic ParA deposition onto the nucleoid like a treadmilling event (**Figs. 1 and 2**). Importantly, the ParA oscillation wave is critically modulated by the nucleoid length and boundary conditions (**Fig. 3**); it is orchestrated by the ParB-mediated negative feedback that is intertwined with the ParA-mediated positive feedback in nucleoid-binding (**Fig. 4**). We suggest that the ParA oscillation, regardless of the see-saw pattern (**Figs. 1 and 2**) or the circulating wave (**Figs. 3 and S3**), may be reminiscent of the underlying feedback between ParA and ParB that ensures the PC partition fidelity.

Our data contradict with a recent work in two ways (42). First, our experiment shows that ParA oscillates anytime over a coherent nucleoid without any discernible sign of constriction (**Fig. 1**), as previously reported (13). Hence, the initiation of ParA oscillation is not coupled to nucleoid splitting. We think that repeated exposure of the cell to blue light (438 nm) for mTurquoise2 imaging may cause phototoxicity and DNA damage response and cell growth arrest and the observed dynamics in the recent work may reflect this unusual condition (**Fig. S2**). Second, our experiment demonstrates that while the PCs are mobile on the nucleoid with local excursions, their movements are largely insensitive to the direction of ParA oscillation wave propagation (**Figs. 1 and S1**). However, the presence of PC is essential for the ParA oscillation (21). Our work suggests that the role of PC in the ParA oscillation is not direct; instead, it controls the level of ParB available for nucleoid-binding and hence the strength of the ParB-mediated negative feedback.

Crucially, we need to take cautions in interpreting our own results as well. Both the PC-bound and the nucleoid-bound ParB_F_ molecules share the same biochemistry in dissociating the ParA_F_ from nucleoid but with different functions. The PC-bound ParB_F_ dissociates the nucleoid-bound ParA to create and follow the ParA_F_ concentration gradient for the directed movements, partitioning, and hence positioning of the PC (7, 9, 10, 14, 15, 17). The nucleoid-bound ParB_F_ controls the overall level of the nucleoid-bound ParA_F_ (**Fig. 4**). Intriguingly, the PC itself regulates the level of nucleoid-bound ParB_F_ by sequestering the majority of the intracellular ParB_F_ (43), thus controlling the fidelity of its own partition. The question is: What is the physical principle underlying this “autonomy” to ensure the fidelity of PC partition? On one hand, the ParB_F_ forms a condensate surrounding the ParS_F_ that defines the PC of the low-copy-number plasmid (15); and both the PC size and the density of the PC-bound ParB_F_ are suggested to critically control the fidelity of the PC partition (15, 44). On the other hand, the PC not only localizes the cytosolic ParA_F_ but manages its partitioning at a critical point in the parameter space, which ensures both the sensitivity and robustness of the process (15). Together, these findings precipitate a series of open questions at the heart of this “autonomy”.

First, at the global level, low-copy-number plasmids encode ParA and ParB genes and autoregulate their own expression level (29, 46, 54). What is molecular logic of this autoregulation that ensures the PC partition fidelity while driving the ParA oscillation?

Second, while the PC-bound ParB controls the level of nucleoid-bound ParA, the PC formation itself depends on the intracellular ParA level. When the ParA level is too high, it will disrupt the formation of the PC (55). With a multitude of ParA-ParB feedback loops, the question is: What underpins the setpoint in partitioning the intracellular ParA_F_ and ParB_F_ between F-plasmids, nucleoid, and cytosol that coordinates the proper PC formation with nucleoid-bound ParA_F_ level for the partition fidelity?

Third, our current work suggests that the ParA_F_ oscillation may be reminiscent of the underlying molecular mechanism to ensure the PC partition fidelity. However, an unanswered question is: Can the ParB_F_-mediated negative feedback still support the fidelity of PC partition without the ParA oscillation? If the answer is yes, then why does the ParA_F_ “need” to oscillate in the first place? After all, the ParA_F_ oscillation will introduce the large cell-to-cell variation in ParA_F_ concentrations (15). This large fluctuation renders the critical-point-operation of PC partition less robust, although the PC-localization of cytosolic ParA_F_ may provide the buffering mechanism and hence ensure the partition fidelity. Nevertheless, wouldn’t the symmetric inheritance of ParA_F_ between the daughter cells work the best in minimizing the noise level? Does the ParA_F_ oscillation along the nucleoid modulate the level of PC-localized cytosolic ParA_F_ and hence more directly impact the fidelity of PC partition? Conversely, if the ParA oscillation is inseparable from the PC partition fidelity, then what exactly is the function of ParA_F_ oscillation? Given that the ParA oscillation occurs only when the nucleoid becomes long enough, does this size-dependence carry any functional role? According to the conventional wisdom, plasmid partition does not tightly couple to the cell cycle of the host cell (53). However, this notion has never been rigorously tested. Can PCs use the ParA oscillation to “survey” the nucleoid length so that they can better synchronize their partition with the timing of cell division? Intriguingly, this survey of nucleoid landscape can be ultrafast (**Fig. 2**), because the ParA see-saw oscillation provides a “highway” that can outrun the diffusional drift of the protein along the nucleoid.

Last, conventional views consider the plasmid and the nucleoid as two separate entities. Interestingly, ParA oscillation seems to be a common feature between the low-copy-number plasmid partition and the bacterial chromosome segregation. What drives the ParA oscillation in the bacterial chromosome segregation? Is there any difference from the plasmid counterpart?

In sum, the ParAB*S* system is an ancient molecular machinery of DNA partition. It is the ancestor or close relative for many intracellular partition and positioning processes in prokaryotes, including bacteria and archaea (31, 56). Understanding the mechanism of ParAB*S*-mediated partition has a wide-range implications in the cellular functions. Despite its tripartite simplicity, the ParAB*S* system survives the evolution with the robust feedback between ParA and ParB at multiple levels that defy the dissection. While our work along with others’ unravel the intriguing excitable dynamics of ParABS system, it begs the deeper questions for the inner working of this molecular machinery. Addressing these questions will be the focus of our future endeavor.

## Materials and Methods

### Bacterial strains and plasmids

All strains are derivatives of *E. coli* K12 and are listed, together with plasmids in **Table S1**. DLT2202 was a spontaneous mutant resistant to streptomycin. The concentrations of antibiotics used for selective bacterial growth were 10 µg.ml^-1^ of chloramphenicol (Cm) and 20 µg.ml^-1^ of ampicillin (Ap). A22 was added at 20 µg.ml^-1^ in exponential growing cultures (OD ∼ 0.3).

All mini-F plasmids are derivatives of pDAG114 (55) and were introduced into recipient strains by CaCl₂-transformation.

### Epifluorescence microscopy

Strains expressing fluorescently tagged proteins were grown overnight at 30°C in M9 minimal medium supplemented with glucose and casamino acids (M9-glucose-CSA). Cultures were diluted 1:250 in fresh medium and incubated at 30°C until OD600nm ∼0.3. Cells were then spotted (0.6 μl) onto 1% agarose pads buffered with M9 1X or M9-glucose-CSA supplemented with A22 (20 µg.ml^-1^).

Fluorescence images were acquired as previously described (57) using an Eclipse TI-E/B wide-field epifluorescence microscope (Nikon) equipped with a phase contrast objective (CFI Plan APO LBDA 100X oil NA 1.45) and Semrock filters for YFP (Ex: 500BP24; DM: 520; Em: 542BP27), CFP (Ex: 438BP24; DM: 458; Em: 483BP32), or mCherry (Ex: 562BP24; DM: 593; Em: 641BP75). Images were captured with an Andor Neo5.5 SCC-02124 camera, using 15-25% illumination from a SpectraX LED light source (Lumencor) and 0.150-0.300 second exposure time depending on the fluorochrome used and the experiment. Image acquisition and analysis were performed using Nis-Elements AR software (Nikon) and ImageJ plugins (MultiStackReg and MultipleKymograph). For clarity in Fig. S3, we performed image denoising with the use of neural networks to display partition complexes using the plugins NIS.ai / denoise.ai of Nis – Elements AR 6.20.00.

### Fluorescence recovery after photobleaching

Fluorescence photobleaching was performed using a Photonic Instrument Micropoint laser system at 488 nm. The cells were imaged once before photobleaching, bleached (65Hz) for 3s in a selected ROI (0.2 x 0.1 µm), and imaged at different intervals (5 to 30s).

### Western immunoblotting

The intracellular level of ParA and ParB was measured from crude cell extracts using the SDS-PAGE system of Invitrogen NuPage Novex Bis Tris Gels (Thermo Fisher Scientific), followed by electrotransfer to nitrocellulose membranes according to the manufacturer’s recommendations (Trans-Blot Turbo Transfer Pack, Biorad). Immunodetection was performed using ECL substrate (Bio-Rad Clarity) with anti-sera raised against affinity-purified ParA or ParB (Eurogentec) using membrane-immobilized samples. Quantifications were performed using ImageLab (BioRad).

## Acknowledgements

We thank Valentin Quèbre for constructing some plasmids and strains used in this study. J.Y.B. and J. L. designed and supervised the research; O.V., T.N., and L.H. performed the computations of the theoretical model; J. R. and C.M.D. performed the experiments. All authors analyzed the data and edited the manuscript. J.Y.B. is supported by a Centre National de la Recherche Scientifique 80Prime grant. T.N. is supported by the National Science Foundation Graduate Research Fellowship. J.L. is supported by Johns Hopkins University Startup Funds, Catalyst Awards, and National Science Foundation grant 2105837.

## Figure Caption

**Figure S1.**
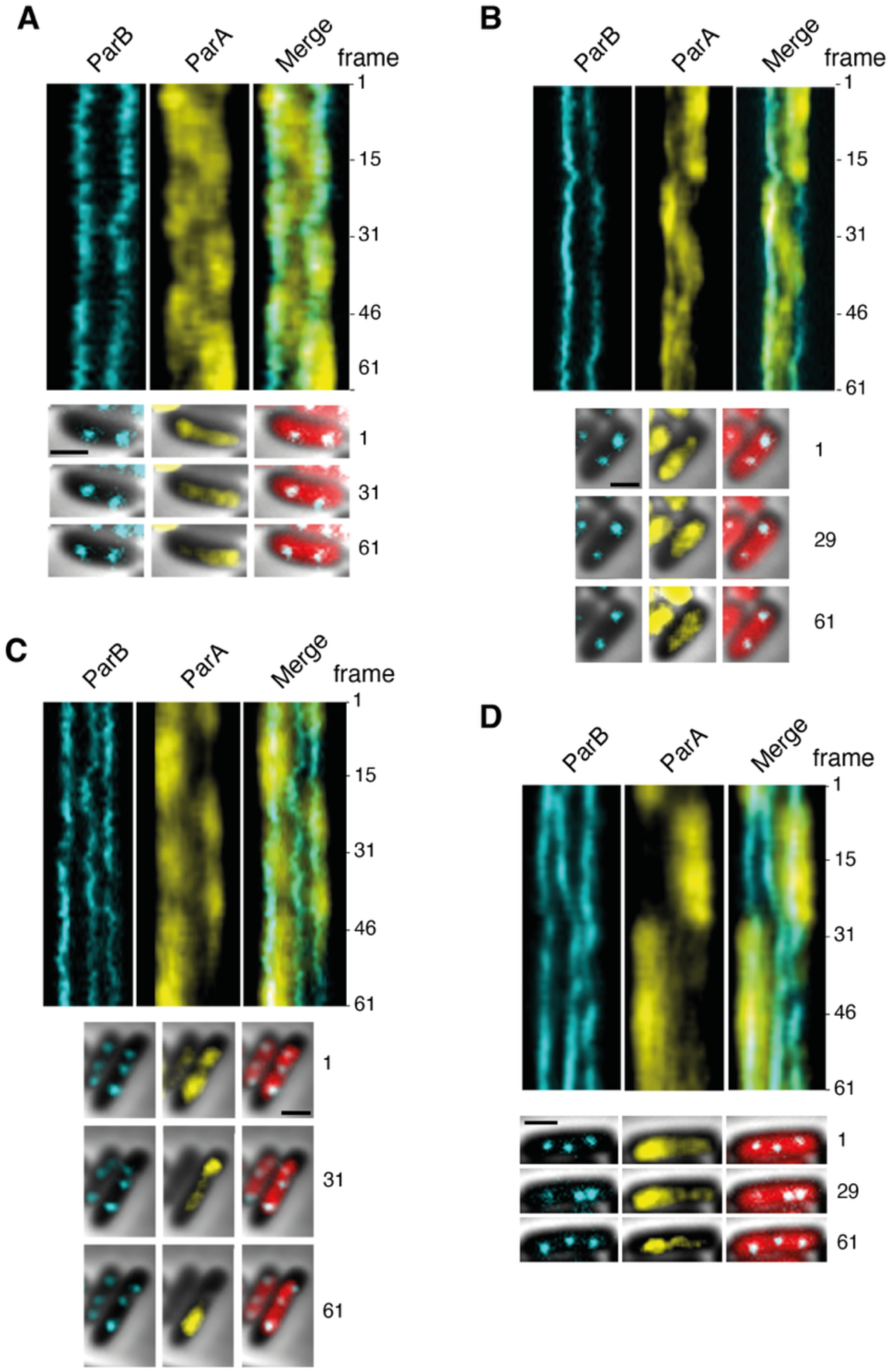
ParA_F_ oscillation does not correlate with directional movement of partition complexes. *E. coli* Hu-mCherry cells (DLT3059), carrying pJYB249 expressing *parA*_F_-mVenus and *parB*_F_-mTurquoise2 were imaged every 30 sec for 30 min from stationary phase cultures. Representative examples display cells with two (A-B) or three (C-D) foci. (*top*) Kymographs in the mTurquoise2 (*left*), mVenus (*central*), and merged mTurquoise2 with mVenus (*right*) channels are shown. Frame number is indicated on the left. (*bottom*) Images of the corresponding cells in the mTurquoise2 (*left*), mVenus (*central*) and merged mTurquoise2 with mCherry (*right*). Note that the cell shown in the panel A is the same as in Fig. 1F. Scale bar, 1 µm.

**Figure S2.**
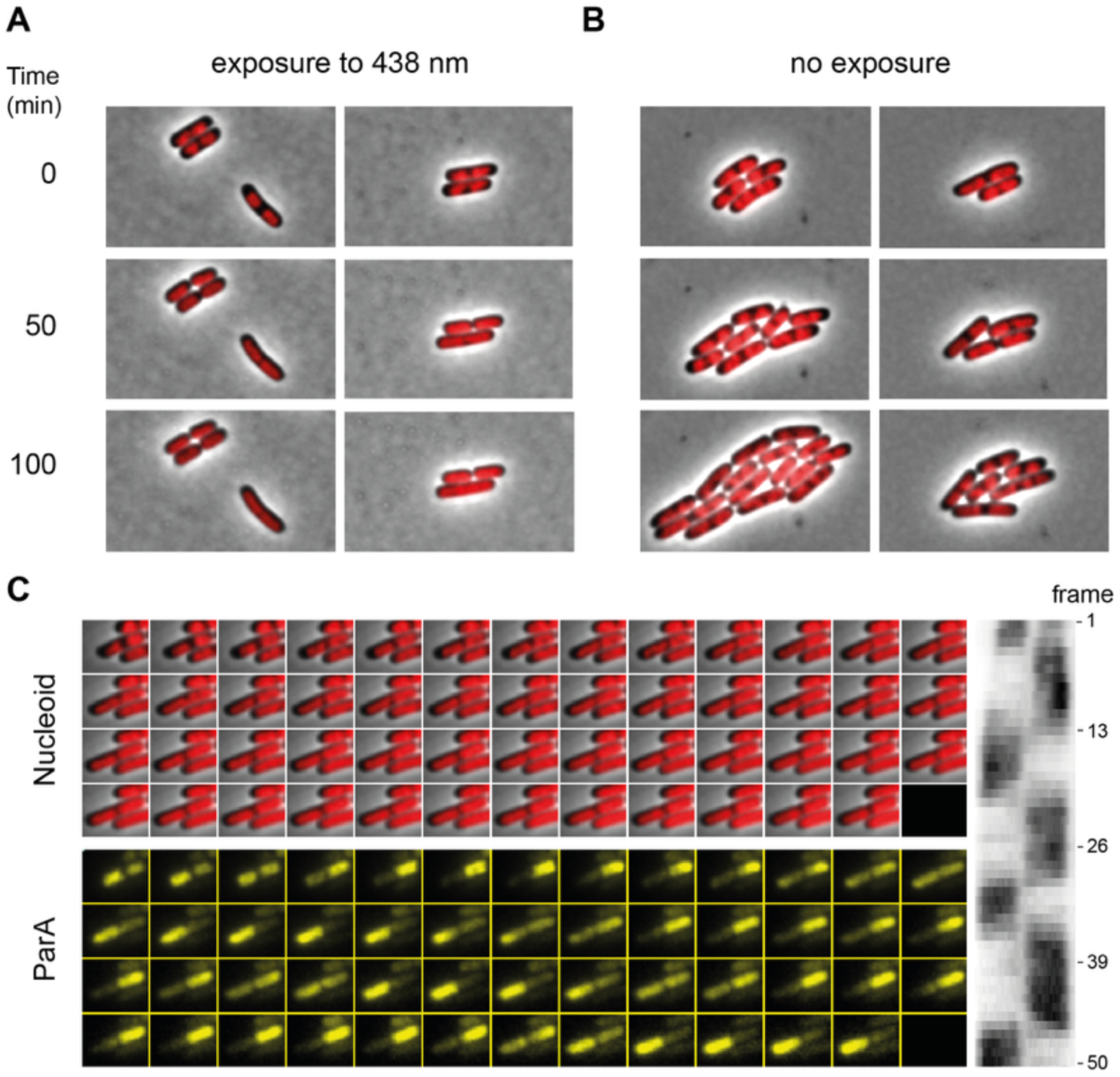
Effect of 438nm exposure to cell growth. An exponentially growing *E. coli* culture was spotted on agarose pad slides and subject A) or not B) to 438 nm exposure every 30 sec. Two representative fields of view are shown for each condition. C) Time-lapse imaging with images taken every 2 min over 100 min in bright field, red, and yellow channels to observe the cell contour, the Hu-mCherry labelled nucleoid, and ParA_F_-mVenus, respectively, and an exposure to the blue channel (438 nm exposure) were added every 30 sec. Under these conditions, cells underwent a growth arrest with a blockage of cell division, and ParA_F_ jumps from one nucleoid to the other. The left panel shows a kymograph from the same cell, displaying ParA_F_-mVenus fluorescence across all frames.

**Figure S3.**
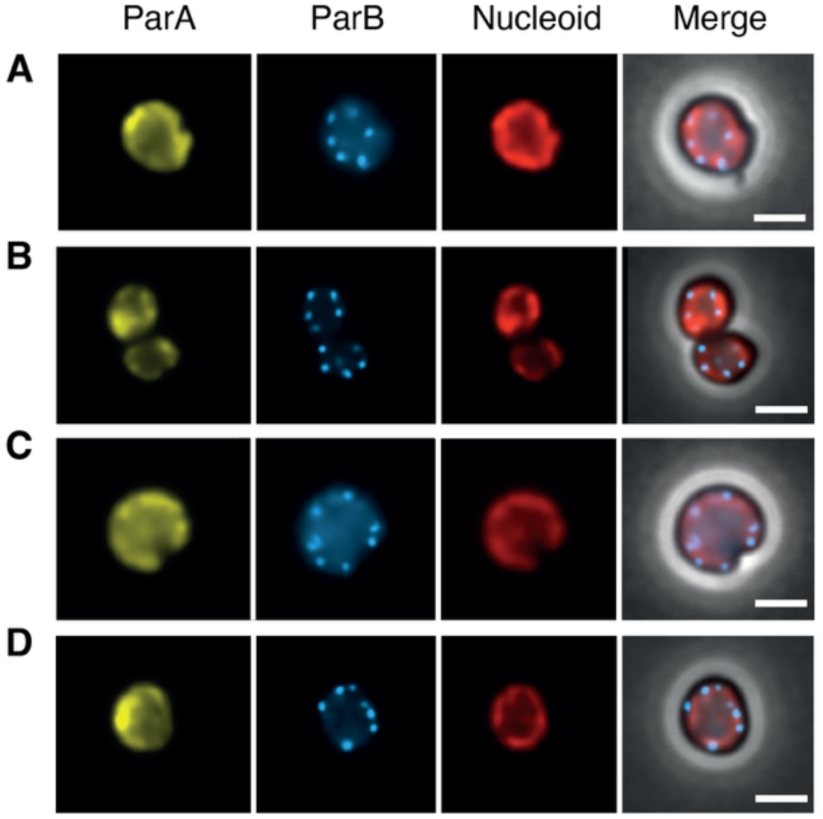
Positioning of partition complexes in A22-treated cells. Representative images of rounded A22-treated cell (DLT3059) displaying donut shaped nucleoids. Bright-field, red, yellow, and cyan channels were acquired sequentially to visualize the cell contour, nucleoid (Hu-mCherry), ParA-mVenus and ParB-mTurquoise2, respectively. Fluorescence images were processed with a denoising step for clarity. Partition complexes appeared regularly spaced along the nucleoids, consistent with functional positioning and active partitioning processes. Scale bar: 2 µm.

**Figure S4.**
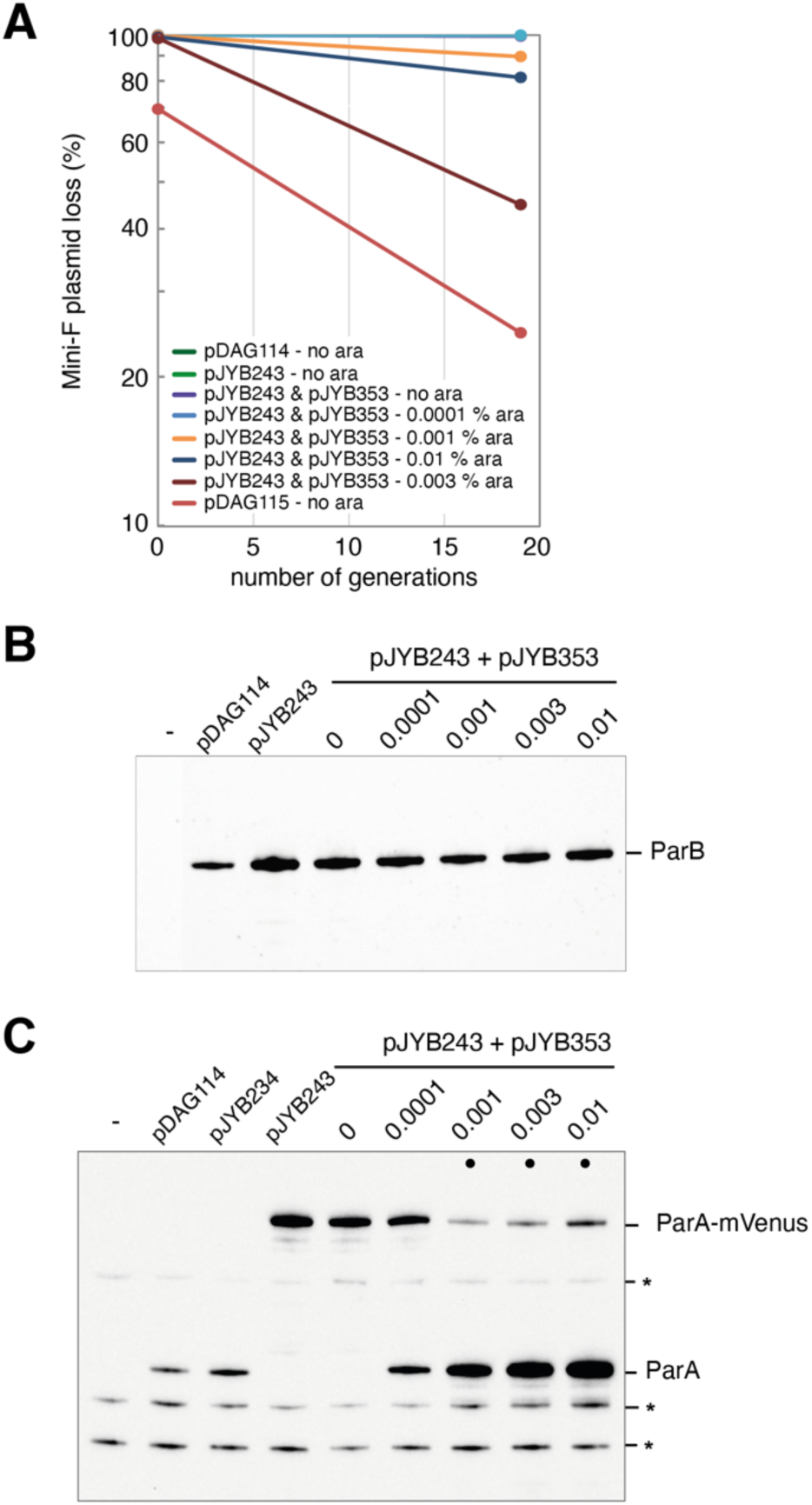
Experiments show that moderate ParA_F_ overexpression preserves both the oscillation and PC partition fidelity, whereas high ParA_F_ overexpression levels disrupt the oscillation and PC partition. All measurements were performed in *E. coli* strain DLT3826 transformed with the indicated plasmid combinations. Plasmids pDAG115 and pJYB243 are derivatives of the wild-type mini-F pDAG114, carrying a *parS* deletion and a *parA*-mVenus fusion, respectively. pJYB353 expresses *parA* under the control of an arabinose-inducible promoter. Arabinose (Ara) was added to the growth medium at the indicated concentrations. A) Western-blot measurements of ParA_F_ levels in the various expression conditions. B) Western-blot measurements of ParB_F_ levels in the same conditions in A). Relative ParA levels representing the total amount of ParA (*i.e.*, ParA and ParA-mVenus combined when both are present), normalized to the level of ParA-mVenus expressed from pJYB243 alone are listed in **Table 2**. C) Plasmid loss rate over generations. Measurements are the average of duplicate experiments, except the data marked with an asterisk (*), which was based on a single replicate.

**Table S1.**
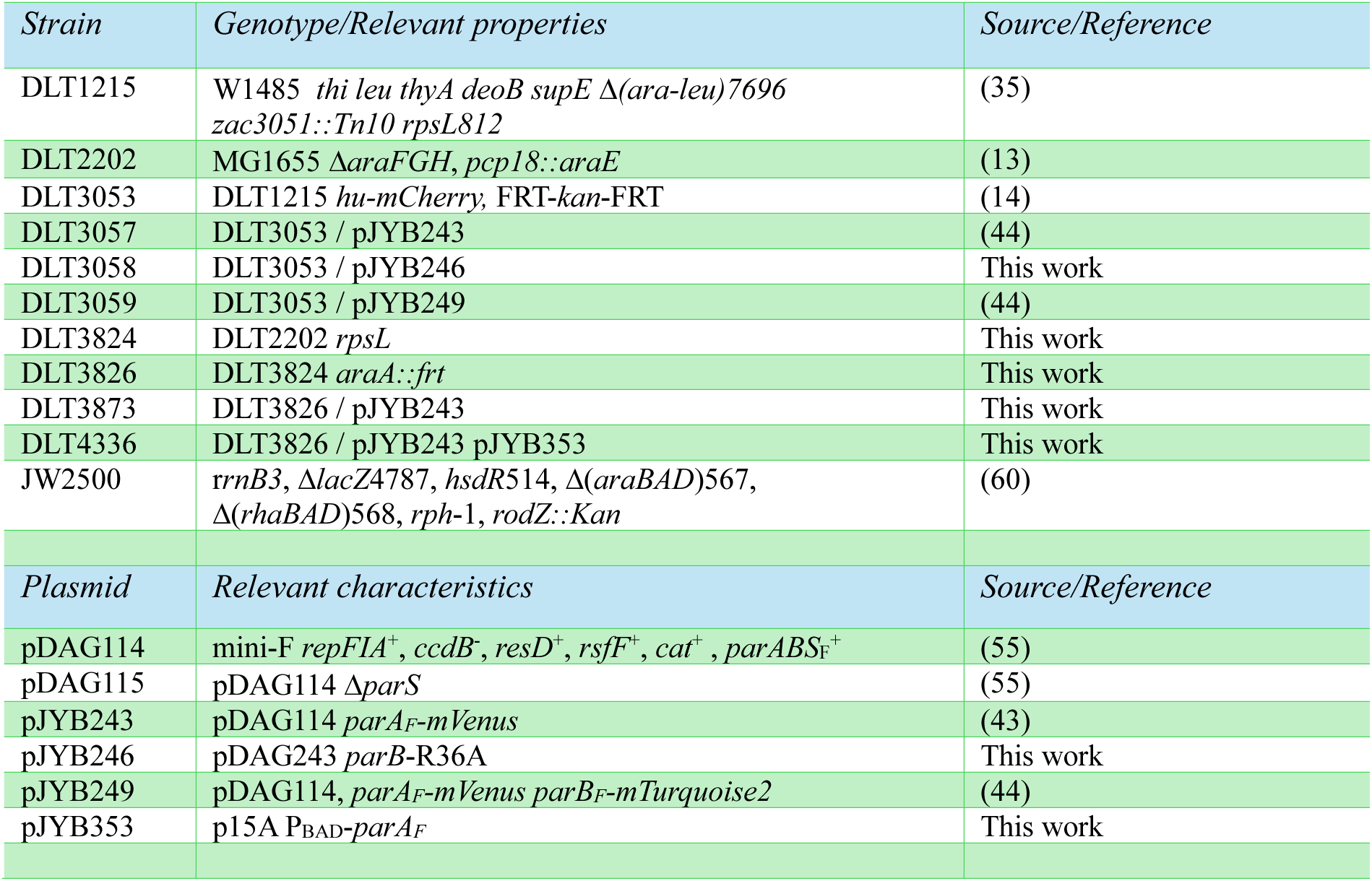
Bacterial strains and plasmids.

